# Cortical Excitation–Inhibition Balance in Autism Varies by Brain Region and Age

**DOI:** 10.64898/2026.06.10.731403

**Authors:** Sergio Osorio, Jasmine Tan, Sheraz Khan, Seppo P Ahlfors, Fahimeh Mamashli, Jussi Alho, Robert M Joseph, Grace Levine, Steven Graham, Gagan Joshi, Zein Nayal, Nicole M McGuiggan, Ainsley Losh, Stephanie Pawlyszyn, Nathaniel Mercaldo, Matti S Hämäläinen, Tal Kenet

## Abstract

Autism spectrum disorder (ASD) is characterized by differences in social communication, interaction, and restricted and repetitive behaviors. The hypothesis that ASD involves altered cortical excitation–inhibition (EI) balance has been extensively investigated, yet evidence remains mixed, partly because EI balance changes substantially during typical maturation. We used resting-state magnetoencephalography (MEG) to examine developmental trajectories of cortical EI alterations with regional specificity in a large cross-sectional cohort (N = 172; 92 typically developing, 80 ASD; ages 6–32). Functional EI (fEI), derived from critical brain dynamics, was estimated across 500 cortical parcellations. Group-averaged fEI maps revealed broadly similar large-scale topographic patterns across groups. Age-stratified analyses revealed increased global fEI in childhood and decreased fEI in adolescence in ASD, with no evidence of a group difference in adulthood. Parcel-wise regressions identified a significant main effect of diagnosis in right dorsolateral prefrontal cortex, where typically developing individuals showed higher fEI across all ages. A significant group-by-age interaction in left inferior parietal cortex indicated diverging developmental trajectories. Finally, fEI in paracentral and precuneus regions was associated with the magnitude of ASD symptoms. These findings indicate that EI imbalance in ASD is neither globally distributed nor static, but expressed through regionally and developmentally specific differences, with relevance for ASD heterogeneity.

## Introduction

Autism spectrum disorder (ASD) is a developmental disorder characterized by differences in social interactions and communication, as well as restricted and repetitive behaviors. It has long been known that the neural correlates associated with ASD are complex and heterogeneous. One prevalent hypothesis has been that ASD is associated with an altered cortical excitation–inhibition (EI) balance (Nelson & Valakh 2015, Port et al 2019, Rubenstein & Merzenich 2003, Sohal & Rubenstein 2019). Converging evidence from genetic, molecular, and circuit-level studies implicates disruptions in neurotransmitter signaling as central contributors to ASD pathophysiology, leading to atypical neuronal excitability and network stability (Blatt et al 2001, Cellot & Cherubini 2014, Drenthen et al 2016, He et al 2021, Kolodny et al 2020, Ma et al 2005, Martin-Ruiz et al 2004, Pizzarelli & Cherubini 2011, Zhao et al 2022). Electrophysiological and neuroimaging studies have further supported this framework, reporting atypical neural variability, oscillatory activity, and functional connectivity patterns in ASD individuals (Ajram et al 2017, Ajram et al 2019, Arutiunian et al 2024, Bruining et al 2020, Carter Leno et al 2022, Manyukhina et al 2022, Orekhova et al 2007, Orekhova et al 2001, Plueckebaum et al 2023). However, despite broad support for an EI imbalance in ASD, results of investigations into this hypothesis remain heterogeneous.

A partial explanation of the observed heterogeneity in results may stem from the fact that in typical development, cortical EI balance is not static across the lifespan but undergoes substantial refinement throughout childhood and adolescence as inhibitory circuitry matures and large-scale networks become increasingly specialized (Caballero et al 2021, Di Cristo 2007, Hensch 2005, Larsen et al 2022, Perica et al 2022, Zhang et al 2024). Developmental processes such as synaptic pruning, myelination, and the progressive strengthening of GABAergic inhibition all contribute to age-related changes in neural variability and network stability, particularly within association cortices (Caballero et al 2021, Sydnor et al 2021, Topchiy et al 2024). Regions within the prefrontal and parietal cortex exhibit especially protracted developmental trajectories, paralleling the maturation of higher-order cognitive and executive functions (Larsen et al 2022 2017, Larsen & Luna 2018 2017, Luna et al 2015 2017, Perica et al 2022 2017, Sydnor et al 2021, Tervo-Clemmens et al 2023). In this context, ASD has been proposed to involve atypical developmental tuning of EI balance rather than a fixed deviation from typical cortical dynamics (Carter Leno et al 2022, LeBlanc & Fagiolini 2011, Nelson & Valakh 2015, Plueckebaum et al 2023). Indeed, these data are supported by studies looking at GABA concentrations in ASD using non-invasive techniques such as magnetic resonance spectroscopy (MRS) and PET, and findings of group differences in childhood, but not in adulthood (Ajram et al 2019, Fung et al 2021). Such altered trajectories could lead to age-dependent and region-specific differences in cortical excitability and network organization, underscoring the importance of understanding developmental effects when examining EI balance in neurodevelopmental disorders.

Noninvasive neurophysiological recordings, specifically millisecond time-resolution EEG and MEG, provide a powerful framework for probing EI balance at the systems level in the human brain, albeit indirectly. For instance, gamma band oscillations indirectly reflect inhibitory interneuron activity, as they arise from interactions between GABAergic fast-spiking interneurons and excitatory pyramidal cells (Buzsaki & Wang 2012, Muthukumaraswamy et al 2015). Other indirect EEG/MEG measures linked to EI balance include the aperiodic (1/f) exponent of the power spectrum, whose slope reflects the relative dominance of excitatory versus inhibitory synaptic currents across large neural populations (Ahmad et al 2022, Donoghue et al 2020). Using EEG and MEG, previous studies have indeed found support for the hypothesis postulating differences in cortical EI ratios in ASD compared to typically developing (TD) individuals, both at rest and during tasks (Arutiunian et al 2024, Bruining et al 2020, Carter Leno et al 2022, Manyukhina et al 2022, Orekhova et al 2007). That said, such studies, to date, have been confined to children, with heterogeneous results across studies, populations, and task designs. Furthermore, non-invasive EEG/MEG studies of EI balance that are limited to the sensor level lack spatial specificity and thus cannot reliably assess region-specific differences in ASD.

Rather than relying on the spectral properties of a single frequency band or the broadband slope, a recently developed approach, functional Excitation-Inhibition, fEI for short, quantifies the dynamic relationship between narrow-band amplitude and the variability of normalized signal fluctuations across time windows, grounded in computational models of critical brain dynamics (Bruining et al 2020). Its theoretical foundation rests on the observation that near-critical neural networks display a characteristic inverse relationship between amplitude and fluctuation variability: high coupling between these two quantities is indicative of a more inhibition-dominated state, while low coupling reflects relative excitation, yielding a fEI ratio defined as 1 minus the Pearson correlation between them. When excitation dominates, activity cascades propagate more freely and when inhibition dominates, cascades are suppressed. Thus, fEI captures the temporal structure of ongoing frequency dynamics, originally demonstrated in the alpha band (Bruining et al 2020), rather than raw power or spectral slope, making it sensitive to shifts in network state that may not be apparent in static spectral measures. Crucially, because fEI is estimated from resting-state time series without requiring task performance, it is especially well suited to large cohorts spanning a wide age range, as in the present study. Importantly, fEI can also be projected into cortical source space, allowing for spatially resolved assessment of fEI across large-scale cortical networks. As such, fEI in combination with MEG source modelling provides a principled framework for investigating how EI dynamics vary across development, brain anatomy, and diagnostic groups.

In this study, we used resting-state magnetoencephalography (MEG) to examine cortical fEI in a large cohort (N=172) of TD (N=92) and ASD (N=80) individuals, spanning childhood, adolescence, and early adulthood (age range 6–32 years). fEI were estimated in source space MEG data across the cortical mantle using anatomically informed parcellations, allowing us to characterize the spatial distribution of fEI at rest and to assess group- and age-related effects on regional fEI. Using parcel-wise regression and cluster-based analyses, we investigated whether developmental trajectories of fEI differed by diagnostic groups, and identified cortical regions exhibiting significant diagnostic group and group-by-age interaction effects. Finally, we tested whether individual variability in cortical fEI was related to clinically validated measures of ASD symptoms.

## Methods

### Participants

Resting-state MEG data were obtained from 172 participants, 92 TD (mean age = 16.8 years) and 80 ASD individuals (mean age = 17.3 years). The full age distributions per group are shown in figure s1. Non-verbal and verbal IQ were measured for all participants, with the Kaufman Brief Intelligence Test II (KBIT-2, Kaufman & Kaufman 2004) used for 102 of the participants, and the Differential Ability Scale (DAS,Elliott 2007) used for the remaining 70 participants. All participants had full scale IQ ≥ 70.

All ASD individuals had a formal clinical diagnosis of ASD. Diagnosis was verified using Module 3 or 4 of the Autism Diagnostic Observation Schedule, Second Edition, (ADOS-2, Lord et al 2012), administered by trained research personnel with demonstrated inter-rater reliability. ADOS scores were based on the combined Social Affect and Restricted Repetitive Behaviors algorithm scores for Modules 3 and 4 (Hus & Lord, 2014). In most of the ASD participants, the diagnosis was further verified by the Social Communication Questionnaire, Lifetime Version (SCQ Lifetime, Rutter et al 2003), completed by a parent, and or the quantitative Social Responsiveness Scale (SRS, Constantino 2012), child version completed by a parent for participants under age 18, or the self-report form for adult participants. ASD participants who either had a borderline ADOS-2 score, did not meet a cutoff of ≥15 on the SCQ, or had a total *t*-score <59 on the SRS, were further evaluated by an expert clinician (RMJ) to confirm the ASD diagnosis. Only participants with a confirmed ASD clinical diagnosis were included in the ASD group. TD participants were screened for ASD symptoms using the SRS, child version completed by a parent for participants under age 18, or the self-report form for adult participants, and or the SCQ, completed by a parent.

Informed consent was provided by adult participants directly, and by parents or legal guardians for underaged participants, in addition to participants’ assent, when possible, in accordance with the protocols approved by the Institutional Review Board of the Massachusetts General Hospital. A summary of clinically assessed scores is presented in table 1.

**Table 1.** Characterization of participants. *p*-values are shown for independent samples *t*-tests for the difference of means across groups. NVIQ (non-verbal IQ), VIQ (Verbal IQ), ADOS (autism diagnostic observation schedule), SCQ (social communication questionnaire – current), SRS (social responsiveness scale).

|  | TD (27 female) |  |  | ASD (15 female) |  |  | p |
| --- | --- | --- | --- | --- | --- | --- | --- |
|  | N | Mean (SD) | Range | N | Mean (SD) | Range |  |
| **Age** | 92 | 16.8 (5.1) | 7-30 | 80 | 17.3 (5.9) | 6-32 | 0.564 |
| **NVIQ** | 92 | 111 (14.3) | 82-140 | 80 | 107.5 (16.9) | 70-144 | 0.124 |
| **VIQ** | 92 | 109.1 (11.8) | 80-142 | 80 | 104.4 (19.8) | 61-147 | 0.053 |
| **SCQ** | 34 | 3.1 (2.7) | 0-13 | 35 | 14.6 (5.7) | 8-30 | $> 0.001$ |
| **SRS** | 61 | 43 (4.7) | 36-55 | 70 | 75.9 (11.4) | 42-100 | $> 0.001$ |
| **ADOS** | - | - | - | 80 | 11.8 (4.7) | 7-27 | - |

### Experimental paradigm

Resting-state MEG data were collected during a continuous 5-min eyes-open fixation paradigm. Participants were instructed to relax and maintain gaze on a central fixation cross projected onto a rear-projection screen positioned 100 cm in front of the participant.

### MRI data acquisition and processing

High-resolution T1-weighted structural MRI scans were acquired on Siemens 1.5-T or 3-T systems using standard MPRAGE sequences using either a 12-channel or a 32-channel head coil. Cortical reconstruction and volumetric segmentation were performed using FreeSurfer (Dale 1999, Fischl et al 1999a, Fischl et al 1999b).

### MEG data acquisition

MEG recordings were collected inside a magnetically shielded room (IMEDCO, Switzerland) using a 306-channel Elekta Neuromag VectorView system (204 planar gradiometers, 102 magnetometers). Data were sampled at either 600 Hz, 1000 Hz or 3000 Hz after filtering between 0.1 and 200 Hz, 330 Hz or 1000 Hz. Head position was tracked continuously using four HPI coils. The location of the HPI coils, the nasion, preauricular points, and additional points on the scalp surface were digitized using a Fastrack digitizer (Polhemus) system. The digitization information was used to co-register the MEG and MRI data for source modeling analyses. Eye movements and cardiac signals (ECG, VEOG, HEOG) were simultaneously recorded and used to detect heartbeat, saccades and blink artifacts. At the end of each session, empty-room data were recorded to measure instrumentation and environmental noise levels in the MEG system.

### MEG data preprocessing

MEG data were preprocessed using MNE python (Gramfort et al 2013, Gramfort et al 2014). First, MEG signals were visually inspected and segments containing gross noise due to excessive movement were manually marked for exclusion. Bad channels were identified automatically using a power spectral density (PSD)–based z-score procedure. Data were filtered between 0.1 and 120 Hz and channels whose PSD exceeded a threshold of 4 z units were flagged as bad. Additional visual inspection was carried out to determine if other channels that were not automatically flagged for deletion should also be rejected. Next, the data were spatially filtered using the signal space separation (SSS) method (Taulu et al 2004, Taulu & Simola 2006), or the Temporal Signal Space Separation (tSSS, Taulu & Simola 2006) methods when possible. While both SSS and tSSS minimize noise from sources outside the virtual sphere circumscribing the MEG sensors, the latter procedure also corrects for head motion using the continuous head position data described in the previous section. Artifacts resulting from cardiac and oculomotor activity were removed using Independent Component Analysis (Lee et al., 1999). Data were filtered between 1 and 45 Hz and then decomposed into 20 independent components. Components reflecting heartbeat, eye movements and blinks were automatically flagged for rejection, whereas additional bad components were identified and rejected by visual inspection.

### Source Modeling

MEG data from the planar gradiometer sensors were projected onto cortical space using MNE-Python (Gramfort et al 2013). Source estimates were computed on each participant’s native cortical anatomy reconstructed from their T1-weighted MRI using FreeSurfer. The forward model was derived using a single-compartment boundary element model (BEM, Hämäläinen & Sarvas 1989), with inner-skull surface meshes generated from individual MRI scans via the watershed algorithm. Cortical current estimates were obtained using Minimum-Norm Estimate method (MNE, Hämäläinen & Ilmoniemi 1994) with a loose orientation constraint of 0.2 and depth weighting of 0.8 (Lin et al., 2006a, 2006b). To account for environmental and instrumental noise, a noise covariance matrix was estimated from empty-room recordings. This covariance matrix was used to construct the inverse operator (Hämäläinen & Ilmoniemi 1994) with a regularization parameter of 0.1.

### Estimation of fEI in cortical space

fEI (Bruining et al 2020) was estimated across 250 approximately equally sized parcellations per cerebral hemisphere. First, individual anatomies were morphed into the fsaverage cortical surface with the spherical morphing method based on sulcal and gyral patterns (Fischl et al 1999b and Dale 1999). The cortical surface was parcellated using the seven network Global-Local functional atlas (Schaefer et al 2018) and source reconstructed signals were averaged across all vertices within each parcellation. fEI was then estimated following the procedure described by Bruining et al. (2020). For each parcellation, the envelope of the band-passed (8–12 Hz) signal segmented into 8-s windows (2000 samples at 250 Hz) with 50% overlap. Within each window, the mean signal amplitude was computed, and the amplitude-demeaned time series was transformed into a cumulative-sum signal profile. This profile was normalized by the window-specific mean amplitude and subsequently linearly detrended. For each window, the normalized fluctuation function nF(t) was computed as the standard deviation of the amplitude-normalized, linearly detrended time series, thus capturing within-window amplitude variability independently of mean signal level. A Pearson correlation was then computed between the window-wise nF(t) variability and the corresponding window-wise mean amplitudes. Parcel-wise fEI was defined as:

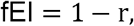

where r denotes the amplitude–nF(t) correlation. Higher fEI values reflect weaker coupling between fluctuation variability and signal amplitude, consistent with a shift toward increased functional excitation relative to inhibition; lower fEI values indicate the converse, reflecting stronger inhibitory engagement. Finally, fEI values were projected back into cortical space with a single value per parcellation for group-effect visualization.

### Computing the effect of group and age in fEI in cortical space

To assess the effects of diagnostic group and age on cortical fEI, we employed Ordinary Least Squares (OLS) linear regression. Parcel-wise fEI values were entered into two linear models to test the effects of age, diagnostic group, and their interaction. An additive model was specified as:

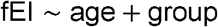

Whereas the interaction model was specified as follows:

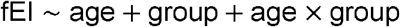

where group was dummy coded with ASD as the reference level (ASD = 0, TD = 1). For each of the 500 parcels, parameter estimates (β coefficients) and p-values for the age effect, the group effect, and the group-by-age interaction were obtained and stored. These parcel-level statistics were mapped back onto the cortical surface by assigning the parcel’s β or p value to all vertices belonging to that parcel, yielding full-brain statistical maps for visualization.

To visually identify spatially contiguous regions exhibiting significant effects on fEI, parcel-wise p values were converted to 1–p and thresholded at p < .05 (uncorrected). Parcels surpassing the threshold were grouped into spatially contiguous clusters. For every subject, mean fEI values within these clusters were computed. These cluster-level fEI values were used for group comparisons, developmental analyses, and visualization.

### Statistical comparison of mean group differences

Group differences in mean hemispheric fEI across age cohorts were assessed using the Mann-Whitney U test, a non-parametric alternative to the independent samples t-test. This approach was selected because global fEI distributions, computed as the mean fEI value across all cortical parcellations per participant, departed from normality, as confirmed by Shapiro-Wilk tests (ASD: W = 0.9679, p = 0.0419; TD: W = 0.9644, p = 0.0129). Separate tests were performed for left and right hemispheres within each age cohort (childhood, adolescence, adulthood). Reported p values are uncorrected. Group differences in continuous demographic and clinical variables (age, IQ, SRS, SCQ) reported in Table 1 were assessed using independent samples t-tests. All tests were two-tailed, and significance was evaluated at an uncorrected threshold of p < 0.05.

## Results

### Group-Level Distribution of Raw fEI

We began the characterization of the spatial distribution of resting-state fEI across the cortex in TD and ASD individuals by computing fEI values in each of the 250 cortical parcellations (Figure 1). Group-averaged surface maps revealed broadly similar large-scale patterns of fEI across the two groups (Figure 1a), for both the left (ASD: mean = 0.54, SD = 0.06; TD: mean = 0.57, SD = 0.06) and right (ASD: mean = 0.54, SD = 0.06; TD: mean = 0.57, SD = 0.06) cortical hemispheres. A histogram of fEI values within each group confirmed this observed trend, and largely overlapped across the two groups (figure 1b).

**Figure 1.**
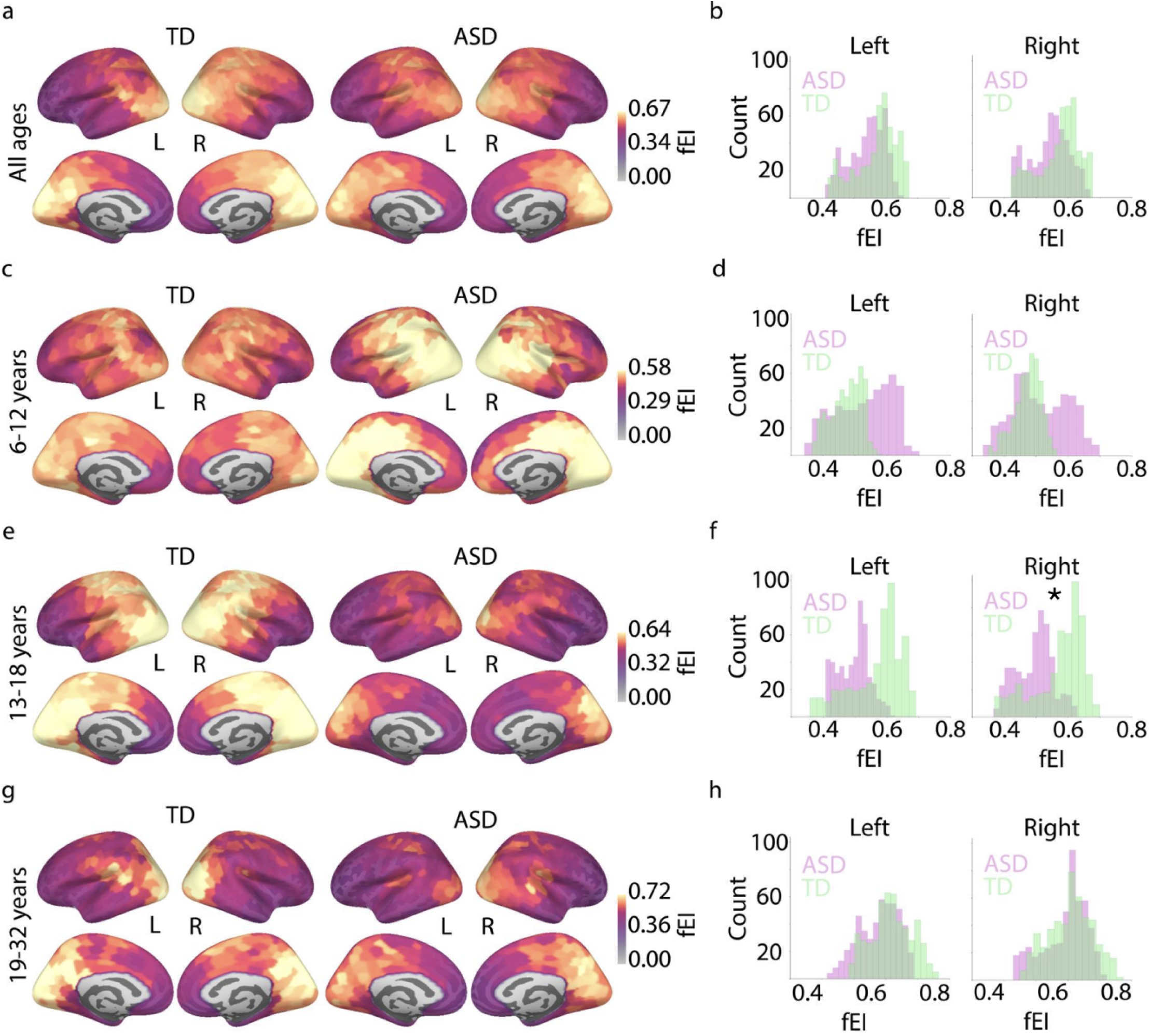
Spatial distribution of raw fEI in TD and ASD groups across childhood, adolescence and adulthood. Cortical distribution of EI ratios across all ages (**a**), between 6 and 12 years (**c**), between 13 and 18 years (**e**), and between 19 and 32 years (**g**). **(b, d, f, h):** Histograms for the EI values in left (left panel) and right (right panel) hemispheres across the four age groupings. Asterisk represents a statistically significant effect (* p < 0.05, uncorrected).

Next, to visualize the potential effect of age, we subdivided our sample into three age groups: childhood (6–12 years), adolescence (13–18 years), and adulthood (19–32 years). Between 6 and 12 years (Figure 1c), more pronounced group differences emerged in the cortical distribution of fEI, with ASD children (N = 18) showing elevated fEI in occipital, parietal, and central regions compared to TD children (N = 16). Histograms of fEI values showed markedly, although not significantly, different means across both left (ASD: mean = 0.53, SD = 0.09; TD: mean = 0.47, SD = 0.05) and right (ASD: mean = 0.51, SD = 0.09; TD: mean = 0.47, SD = 0.04) hemispheres (Figure 1d). Between 13 and 18 years, this pattern reversed, with TD adolescents (N = 41) showing higher fEI in occipital, parietal, and central regions than the ASD subgroup (N = 27; Figure 1e). Histograms confirmed higher mean fEI in TD (left: mean = 0.57, SD = 0.08; right: mean = 0.57, SD = 0.07) compared to ASD (left: mean = 0.50, SD = 0.05; right: mean = 0.50, SD = 0.06) adolescents (Figure 1f). This difference was statistically significant only for the right hemisphere (Mann-Whitney U = 392, p = 0.04, uncorrected). Finally, the cortical distribution of fEI appeared to stabilize between 19 and 32 years, showed occipital and posterior cingulate peaks emerging in both ASD (N = 28) and TD (N = 26) groups (Figure 1g), and similar mean fEI values in left (ASD: mean = 0.58, SD = 0.06; TD: mean = 0.60, SD = 0.06) and right (ASD: mean = 0.59, SD = 0.07; TD: mean = 0.61, SD = 0.07) cortical hemispheres (Figure 1h).

### Main Effect of Diagnostic Group on Cortical fEI

To further characterize the relationship between fEI, diagnostic group, and age, we fitted an additive linear regression model with fEI as the outcome variable and group and age as predictors. Parcel-wise regression analyses revealed greater beta coefficients for the effect of diagnostic group, independent of age, in prefrontal, inferior temporal and inferior parietal regions (figure 2a). Corresponding parcellation-wise significance maps demonstrated that several clusters survived an uncorrected threshold of p < 0.05, indicating statistically reliable group differences, particularly in right prefrontal regions (figure 2b). This prefrontal cluster included six anatomical sub parcellations, located in the region of the dorsolateral prefrontal cortex (DLPFC, mean β = 0.09, CI = [0.08, 0.11], mean p = 0.01 figure 2c and supplementary materials, table s1). Mean fEI values extracted from this right DLPFC cluster showed a positive relationship with age for both ASD (β = 0.008, CI = [0.002, 0.013], p = 0.02) and TD (β = 0.009, CI = [0.001, 0.017], p = 0.03) groups, with older participants showing higher mean fEI values across both groups, as well as an overall group difference in mean fEI across all ages (figure 2d).

**Figure 2.**
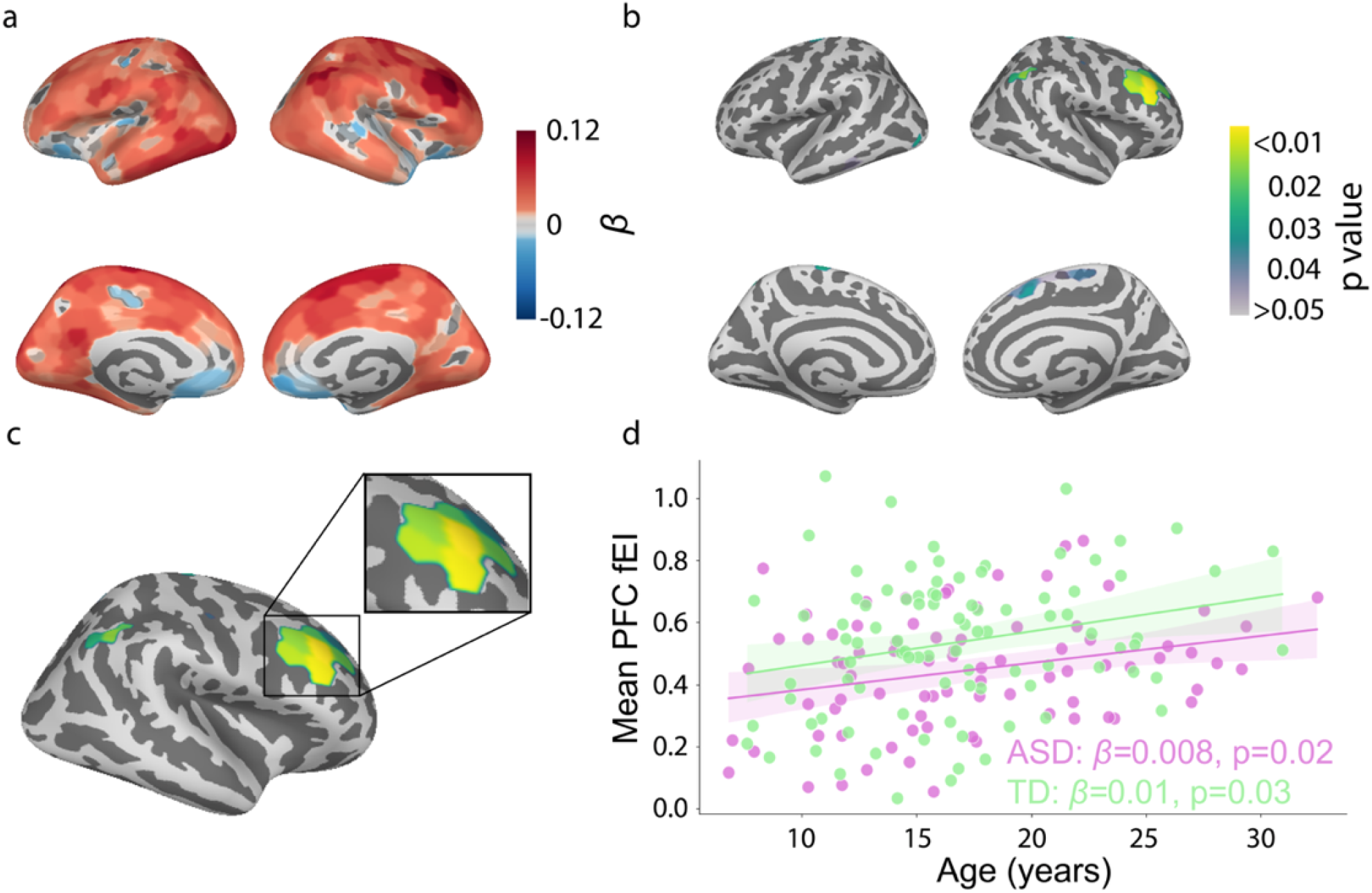
Main effect of group on fEI. Surface-based maps show the results of linear regression. **a**. Regression coefficients (β) for the group contrast projected onto the fsaverage cortical template. Warm colors indicate higher fEI values in the TD group relative to ASD, whereas cool colors indicate higher fEI in ASD relative to TD. **b**. Uncorrected p-value maps for the group effect displayed on the same cortical surfaces. Colored regions indicate sub-parcellations showing statistically significant group differences, thresholded at p < 0.05. **c**. Significant regression coefficients from six anatomical labels were primarily clustered in the right DLPFC. **d**. fEI values extracted from and averaged across these six significant clusters of the right DLPFC as a function of age.

### Diverging developmental trajectories between ASD and TD individuals are expressed in the parietal cortex

To assess whether age-related changes in fEI differed between diagnostic groups, we fitted an additional linear regression model that included the main effects of group and age as well as their interaction. The group × age interaction revealed spatially specific modulation of fEI across the cortex, concentrating in prefrontal, central, parietal and occipital regions (figure 3a). Parcellation-wise significance maps revealed significant clusters at an uncorrected threshold of p < 0.05 primarily located in inferior parietal and inferior central cortical areas (figure 3b). The biggest of these clusters was in the left inferior parietal cortex (IPC) and included six anatomical sub-parcellations (mean β = 0.016, CI = [0.015, 0.017], mean p = 0.02, figure 3c and supplementary materials, table s1). Finally, age was positively associated with parietal fEI values among TD individuals (β = 0.012, CI = [0.003, 0.022], p = 0.006), but not in ASD individuals (β = 0.001, CI = [−0.006, 0.008], p = 0.84, figure 3d).

**Figure 3.**
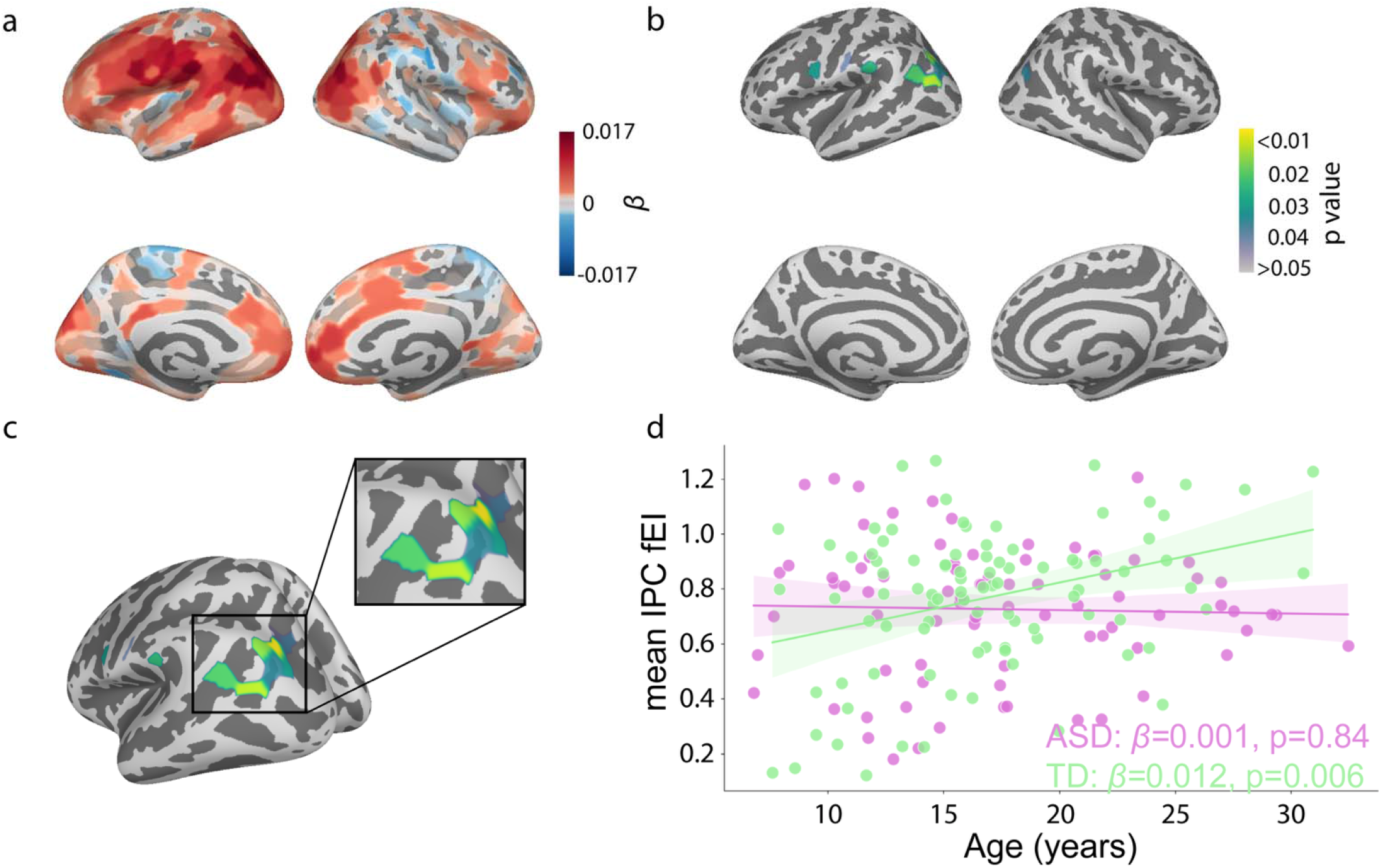
Group × age interaction on cortical fEI. Surface-based maps show the interaction between diagnostic group and age. **a**. Parcellation-wise regression coefficients (β) for the group × age interaction displayed on the fsaverage cortical template. Positive β values indicate regions where age-related changes in fEI are greater in the TD group relative to the ASD group, whereas negative values indicate stronger age-related effects in ASD. **b**. Uncorrected p-value maps for the interaction effect. Colored regions represent vertices showing statistically significant group × age interactions (p < 0.05), while gray regions denote non-significant effects. **c**. Significant values are observed in six sub parcellations, primarily clustered in the left inferior parietal cortex. **d**. fEI values extracted from and averaged across these six significant sub-parcellations, plotted as a function of age.

### Functional excitation–inhibition balance predicts extent of autism symptoms

Lastly, we sought to assess whether cortical fEI was associated with ASD symptoms. We focused on the ADOS algorithm total score as the ASD symptom measure for two reasons. First, it was available for all ASD participants, and second, the SRS was administered in different versions (parent-report vs. self-report) across participants, introducing potential inconsistencies.

We began by testing whether the mean fEI within each of the two clusters (frontal and parietal) that showed significant between-group effects correlated with the total ADOS score, finding no evidence for an association of the ADOS score with the fEI in the DLPFC (r = −0.1, CI = [−0.32, 0.13], p = 0.39) or in the IPC (r = −0.17, CI = [−0.38, 0.07], p = 0.13).

Then, to further estimate whether fEI was associated with ASD symptom severity, we used a parcel-wise linear regression in the ASD group (Figure 4a). Surface-based maps of regression coefficients revealed widespread negative associations between fEI and ADOS scores, independent of age, predominantly in medial, central, parietal and occipital regions, suggesting that higher fEI was associated with lower ADOS scores, i.e. less pronounced ASD symptoms. Conversely, prefrontal and orbitofrontal regions showed a positive relationship between fEI and ADOS scores. Corresponding p-value maps, thresholded at an uncorrected p < 0.05 for exploratory visualization, identified spatially specific clusters showing statistically significant associations between fEI and extent of ASD symptoms, primarily located in paracentral and precuneus cortical regions (Figure 4b).

**Figure 4.**
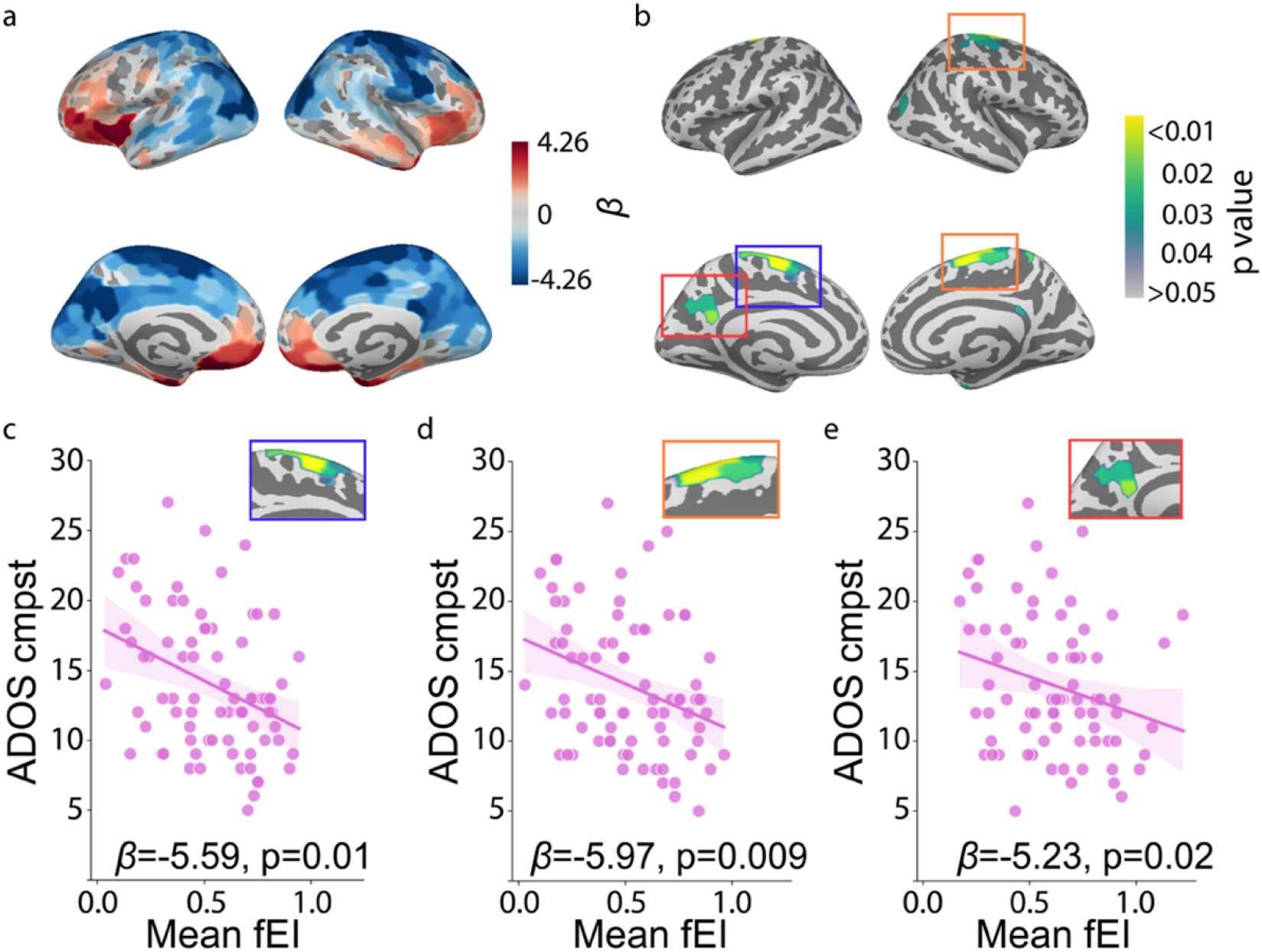
Functional excitation–inhibition balance is associated with autism symptom severity. **a**. Surface-based maps showing regression coefficients (β) for the main effect of fEI ratio in ADOS composite scores, after controlling for age. Negative β values indicate regions where higher fEI is associated with lower ADOS scores. **b**. P-value maps for the main effect of fEI, thresholded at an uncorrected p < 0.05 for visualization. Colored regions indicate cortical areas showing statistically significant associations between fEI and ADOS scores. Gray regions denote non-significant effects. **c**. Relationship between mean fEI extracted from the left paracentral cortical cluster (blue box) and ADOS scores across ASD participants. **d**. Relationship between mean fEI extracted from the right paracentral cluster (orange box) and ADOS scores. **e**. Relationship between mean fEI extracted from the precuneus (red box) and ADOS scores. Reported β and p values correspond to the mean values across all parcellations within the cluster. For the individual beta and p values within each anatomical parcellation see supplementary materials, table s2.

To characterize these effects at the regional level, mean fEI values were extracted from the significant paracentral and precuneus clusters shown in Figure 4b. Across clusters, higher mean fEI within these clusters was associated with lower ADOS composite scores (left paracentral: β = −5.59, CI = [−5.96, −5.22], p = 0.01; right paracentral: β = −5.97, CI = [−6.41, −5.51], p = 0.009; left precuneus: β = −5.23, CI = [−5.63, −4.99], p = 0.02), indicating that more inhibition-dominated fEI values in these regions corresponded to greater ASD symptom severity, independently of age (Figures 4c-e).

## Discussion

This study used resting-state MEG source imaging to characterize the spatial distribution and developmental trajectory of functional excitation–inhibition (fEI) balance in a large cohort of ASD and TD individuals spanning childhood through early adulthood. We found both age-dependent and age-independent differences in fEI between the two groups, and these differences were regionally concentrated rather than globally distributed across the cortex. Age-independent group differences localized to the right DLPFC, while age specific differences in childhood and adolescence localized to posterior regions of the cortex, where an age by group interaction revealed a divergence of developmental trajectories of fEI between the ASD and TD groups in the left inferior parietal cortex (IPC). Lastly, higher fEI in paracentral and precuneus cortical areas was associated with less pronounced ASD symptoms. Together, these findings indicate that fEI differences in ASD are regionally and developmentally heterogeneous, and more locally, can manifest as both a developmentally dynamic divergence (left IPC) or a persistent, age-independent reduction (right DLPFC).

### Group Differences in fEI Show Age Cohort Specific Patterns

When using age as a discrete divider rather than continuous variable, and breaking up the cohort into three distinct groups by age range, the most striking feature was an apparent shift in the ASD group, from elevated fEI in occipito-parietal regions in childhood, to reduced fEI in these same regions during adolescence, and finally apparent equalizing of fEI with the TD group in adulthood. The elevated fEI in childhood aligns with a sensor-level EEG study reporting globally elevated fEI in children with ASD (Bruining et al 2020, Manyukhina et al 2022). It also aligns with studies of children with ASD showing an elevated EI balance (Ajram et al 2019, Bejjani et al 2012, Di et al 2020, Ford et al 2020, He et al 2021, Keehn et al 2017, Kolodny et al 2020, Oliveira et al 2018, Orekhova et al 2019, Siegel-Ramsay et al 2021), as well as with studies showing divergent patterns of functional connectivity differences between children and adolescents with ASD (Dajani & Uddin 2015, Mamashli et al 2018, Vakorin et al 2017, Wiggins et al 2011). Perhaps most importantly, the difference in patterns by age cohorts may explain some of the discrepancies across studies to date, especially given that many studies of ASD routinely include age cohorts that span both childhood and adolescence.

### A Global Shift in fEI in the Prefrontal Cortex

ASD individuals showed significantly lower fEI than TD individuals across all ages in the right DLPFC. This aligns with frameworks proposing that EI alterations in autism are region-dependent rather than pan-cortical (Nelson & Valakh 2015, Port et al 2019). The localization to DLPFC is consistent with post-mortem studies demonstrating reduced parvalbumin-expressing interneuron density specifically in prefrontal areas (BA9, BA46, BA47) in ASD individuals (Hashemi et al 2018), which would shift the local EI ratio toward relative excitation and attenuate the fEI metric by weakening amplitude-fluctuation coupling. Furthermore, preclinical work in several ASD mouse models have shown convergent evidence of disrupted prefrontal inhibitory tone, including altered GABAergic drive and impaired parvalbumin interneuron function (Caballero et al 2021).

The significance of the prefrontal effect is reinforced by the complementary neuroimaging literature. Using MR spectroscopy in a large longitudinal cohort, McKeon et al. demonstrated that adolescent maturation of prefrontal EI balance, reflected in rising glutamate:GABA ratios, supports the development of working memory and executive function (McKeon et al 2024). The present finding that DLPFC fEI is lower in ASD across development may therefore reflect a circuit-level substrate for the executive function difficulties commonly observed in this population (Braden et al 2017, Demetriou et al 2019, Li et al 2024, Robinson et al 2009, Rosenthal et al 2013). Of note, the group difference survived the inclusion of age as a covariate, indicating that this prefrontal deficit is not attributable to developmental delay, but instead represents a persistent deviation in the functional balance of local excitatory and inhibitory processes.

The right DLPFC has also been consistently implicated in studies of ASD, as showing differences in functional connectivity and activation in ASD versus TD individuals, by our group (Osorio et al 2025) and others (Choi et al 2015, Delmonte et al 2013, Doyle-Thomas et al 2013, Hou et al 2022, Richey et al 2022).

### Diverging Developmental Trajectories in Inferior Parietal Cortex

In TD individuals, parietal fEI increased steadily with age, consistent with the progressive maturation of GABAergic inhibitory tone and the refinement of local cortical circuits across adolescence (Sydnor et al 2021, Topchiy et al 2024). In contrast, ASD individuals showed a flat or absent age-related increase in parietal fEI, suggesting a failure to undergo the typical developmental strengthening of inhibitory regulation in this region. These data extend prior work reporting that the developmental trajectory of fEI in children with ASD was altered relative to non-ASD peers, and was associated with language abilities (Plueckebaum et al 2023). It also aligned with prior findings by our group (Alho et al 2021, Khan et al 2018, Kitzbichler et al 2015, Mamashli et al 2018, Mamashli et al 2021) and others (Alaerts et al 2015, Edgar et al 2019, Edgar et al 2016, Fossum et al 2021, Gage et al 2003, Green et al 2022, Green et al 2023, Luna et al 2007, Murphy et al 2017, Port et al 2016, Vakorin et al 2017, Vogan et al 2019, Washington et al 2013), showing different maturational trajectories of brain function and structure among ASD individuals compared to typically developing controls.

A potential additional significance of these findings is derived from the fact that the inferior parietal lobule is among the latest-maturing cortical regions in the human brain, exhibiting protracted structural and functional development well into the third decade of life (Sydnor et al 2021). It supports a broad range of higher-order cognitive operations including attention, social cognition, and multisensory integration (Caspers et al 2013), all of which are implicated in ASD. A failure to develop typical levels of inhibitory engagement in the parietal cortex could result in degraded signal-to-noise ratio in parietal networks, impairing their capacity to filter task-relevant from task-irrelevant information during development. This interpretation is consistent with the broader EI imbalance framework in ASD, wherein reduced inhibitory gain disrupts the precision of cortical representations (Nelson & Valakh 2015, Rubenstein & Merzenich 2003, Sohal & Rubenstein 2019), and aligns with the observation that association cortices represent a nexus of neurodevelopmental risk across psychiatric disorders (Sydnor et al 2021).

### Cortical EI Balance and ASD Symptoms

Within the ASD group, higher fEI in paracentral and precuneus cortical regions was associated with lower ADOS composite scores, indicating that individuals with more TD-like (higher) fEI in these areas had milder symptom profiles. Some of the regions identified partially overlap with the default mode network (DMN), particularly the precuneus. These DMN nodes are critically involved in social cognition, self-referential processing, and mentalizing, functions that are specifically impaired in ASD (Assaf et al 2010). The negative relationship between fEI and ADOS scores suggests that lower EI ratios in these regions, reflecting a relative shift toward inhibition or disrupted amplitude–fluctuation coupling, may correspond to greater disruptions in the neural processes supporting social behavior.

Converging evidence from resting-state fMRI has demonstrated that DMN connectivity is among the most consistently disrupted network features in ASD (Duan et al 2023, Monk et al 2009, Padmanabhan et al 2017), and that the degree of disruption correlates with social symptom severity as measured by the ADOS and SRS (Assaf et al 2010, Jann et al 2015, Yerys et al 2015). The current MEG-based findings extend this literature by showing that the local EI dynamics within DMN nodes themselves, rather than just their connectivity patterns, relate to symptom burden. This is conceptually consistent with the view that EI imbalance in key cortical nodes propagates to disrupt network-level information routing, ultimately manifesting as behavioral deficits (Bruining et al 2020, Sohal & Rubenstein 2019).

### Limitations

The cross-sectional design precludes causal inference about developmental trajectories, and the diverging age-related patterns observed in the parietal cluster should be interpreted as cross-sectional age effects rather than intra-individual longitudinal change. Longitudinal replication would be necessary to confirm whether the apparent failure of parietal fEI maturation in ASD reflects a true within-person trajectory difference. Additionally, while MEG source reconstruction allows spatially resolved imaging of EI balance, source modeling introduces spatial smoothing and potential leakage between adjacent parcellations. The use of individual MRI-based anatomy partially mitigates this concern, but some blurring of anatomically distinct EI effects remains possible. It is also worth noting that the cohort sizes for the district within age range (childhood, adolescence, adulthood) analyses were much smaller than the total cohort size. Another limitation is that participants included in this analysis were recruited across multiple studies, and did not all undergo the same behavioral battery. In particular, SRS and SCQ scores were not available for the full sample, limiting the power of behavioral correlation analyses. Future prospective cohort studies with a uniform behavioral protocol would be better positioned to characterize fEI–behavior relationships across a wider range of ASD phenotypes.

## Conclusions

This study provides the first spatially resolved, MEG-based characterization of fEI balance across the cortical mantle in a large developmental cohort of ASD and TD individuals. The results demonstrate that EI imbalance in ASD, as measured using fEI, is regionally concentrated in the prefrontal and parietal association cortices, with distinct developmental dynamics in each. Age dependence was apparent in the left IPC, where fEI increased with age in the TD group but not the ASD group, and also in broader occipito-parietal areas, where children and adolescents with ASD showed opposite trends relative to TD individuals. In contrast, in the DLPFC, fEI was consistently lower in ASD, irrespective of age. Finally, the association between fEI and ASD symptoms in paracentral and precuneus cortical regions points toward a potential neurophysiological correlate of ASD phenotypic heterogeneity. In combination, these observations may serve as explanatory factors not just of ASD symptoms, but also for some of the heterogeneity observed across ASD studies targeting the EI imbalance hypothesis.

## Conflict of interest

The authors declare no conflict of interest.

## Notes

### Competing Interest Statement

The authors have declared no competing interest.

### Funding

- NIH: P41EB030006, P41EB015896, R01MH117998, S10OD030469, R01DC016765

