## Supplement for "Cortical Excitation–Inhibition Balance in Autism Varies by Brain Region and Age"

**Contents:**

Figure s1

Table s1

Table s2

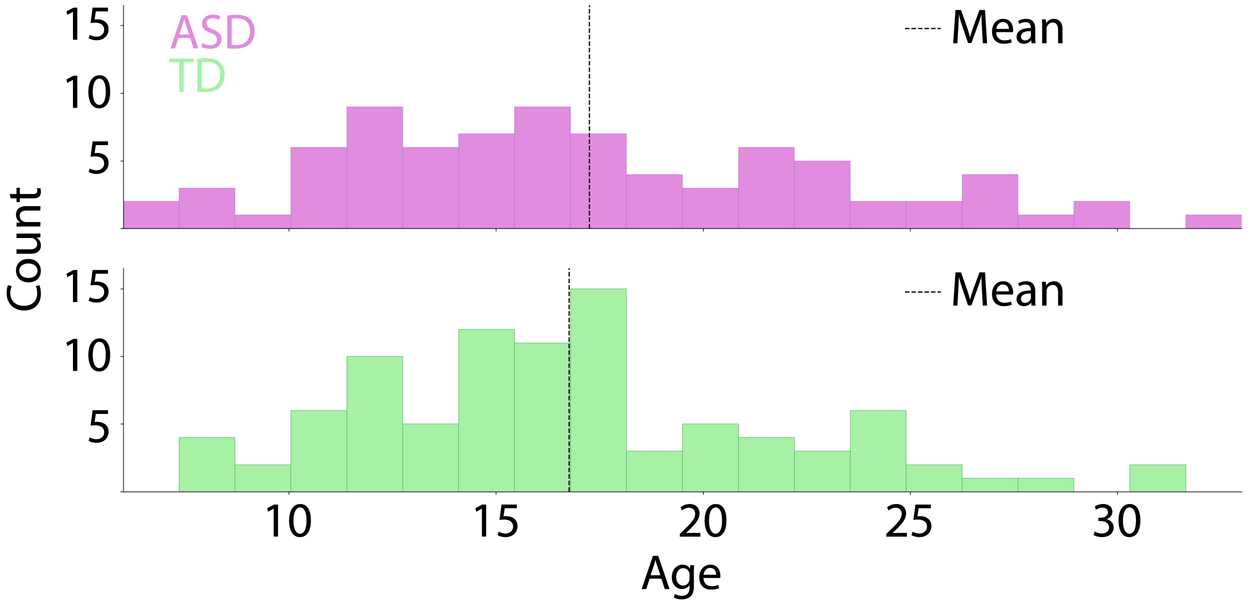


**Figure s1.** Age distribution for the ASD (mean = 17.3 years, SD = 6 years) and TD (mean = 16.8 years, SD 5.1 years) groups. Mean age is illustrated by dotted lines intersecting the x axis.

| Cluster | Label | beta | pval |
|---|---|---|---|
| **Prefrontal** | 7Networks_RH_Cont_PFCl_12-rh | 0.10 | 0.002 |
| | 7Networks_RH_Cont_PFCl_15-rh | 0.12 | 0.000 |
| | 7Networks_RH_Cont_PFCl_16-rh | 0.10 | 0.009 |
| | 7Networks_RH_Cont_PFCl_17-rh | 0.09 | 0.014 |
| | 7Networks_RH_Default_PFCdPFCm_11-rh | 0.07 | 0.031 |
| | 7Networks_RH_SalVentAttn_PFCl_1-rh | 0.08 | 0.012 |
| **Parietal** | 7Networks_LH_Cont_Par_1-lh | 0.02 | 0.007 |
| | 7Networks_LH_Default_Par_2-lh | 0.02 | 0.013 |
| | 7Networks_LH_Default_Par_4-lh | 0.02 | 0.020 |
| | 7Networks_LH_Default_Par_5-lh | 0.02 | 0.033 |
| | 7Networks_LH_DorsAttn_Post_5-lh | 0.01 | 0.036 |
| | 7Networks_LH_Vis_36-lh | 0.02 | 0.028 |

**Table s1.** Individual beta coefficients and p values for each of the anatomical parcellations that make up each of the two clusters where a significant effect was found for group (prefrontal) and the group x age interaction (parietal).

| Cluster | Label | beta | pval |
|---|---|---|---|
| **RH Dorsomedial** | 7Networks_RH_SalVentAttn_Med_12-rh | -6.10 | 0.005 |
| | 7Networks_RH_SalVentAttn_Med_13-rh | -5.47 | 0.017 |
| | 7Networks_RH_SomMot_36-rh | -5.36 | 0.019 |
| | 7Networks_RH_SomMot_40-rh | -7.01 | 0.002 |
| | 7Networks_RH_SomMot_43-rh | -5.95 | 0.005 |
| | 7Networks_RH_SomMot_45-rh | -6.50 | 0.003 |
| | 7Networks_RH_SomMot_48-rh | -6.46 | 0.004 |
| | 7Networks_RH_SomMot_49-rh | -4.91 | 0.018 |
| **LH Dorsomedial** | 7Networks_LH_SalVentAttn_Med_10-lh | -5.24 | 0.016 |
| | 7Networks_LH_SomMot_34-lh | -6.03 | 0.003 |
| | 7Networks_LH_SomMot_42-lh | -6.08 | 0.006 |
| | 7Networks_LH_SomMot_44-lh | -5.02 | 0.014 |
| | 7Networks_LH_SomMot_45-lh | -5.56 | 0.013 |
| **LH PCC** | 7Networks_LH_Default_pCunPCC_4-lh | -5.63 | 0.010 |
| | 7Networks_LH_Default_pCunPCC_6-lh | -4.99 | 0.022 |
| | 7Networks_LH_Default_pCunPCC_7-lh | -5.07 | 0.023 |

**Table s2.** Individual beta coefficients and p values for each of the anatomical parcellations that make up each of the three clusters where fEI significantly predicts ADOS scores.
